# Classification of indeterminate and off-target cell types within human kidney organoid differentiation

**DOI:** 10.1101/2025.05.16.654519

**Authors:** Sean B. Wilson, Jessica M. Vanslambrouck, Angela Murphy, Drew R. Neavin, Joseph E. Powell, Sara E. Howden, Melissa H. Little

**Affiliations:** Novo Nordisk Foundation Centre for Stem Cell Medicine, Murdoch Children’s Research Institute, Parkville, Melbourne, 3052, Australia; Novo Nordisk Foundation Centre for Stem Cell Medicine, Faculty of Health and Medical Sciences, University of Copenhagen, DK-2200 Copenhagen, Denmark; Department of Paediatrics, The University of Melbourne, VIC, Australia; Garvan Institute of Medical Research, Sydney, 2010, NSW, Australia; UNSW Cellular Genomics Futures Institute, University of New South Wales, Sydney, 2052, NSW, Australia

**Keywords:** directed differentiation, kidney organoid, cellular identity, transcriptional profiling, asynchronous differentiation

## Abstract

Human pluripotent stem cell-derived organoids are multicellular models of developing tissues proposed to recapitulate developmental stages of lineage commitment across time. Assessing how accurately protocols recapitulate development is challenged by the lack of accurate reference data sets for early human development, the presence of ‘off-target’ states and the potential to form cellular states not present *in vivo*. This study addresses these challenges with respect to differentiation of human pluripotent stem cells to kidney organoids. Based on a factorial single-cell transcriptomic analysis across a 27-day differentiation protocol, including 150,957 cells collected at five time points, we present a comprehensive classification of predicted identity, representing an *in vitro* temporal transcriptional atlas. For early stages, cellular identity was defined in relation to existing human developmental data across time and organ system. As renal structures arose, this was coupled with the kidney-specific classification methods. In this way we identify predicted cellular states, predictable off-target cell types based on embryology and transitional mesodermal populations with no clear *in vivo* equivalent. An analysis of all existing single cell data on this same protocol suggests the presence of the same off-target trajectories irrespective of cell line or laboratory. This study provides a valuable resource to better identify and understand how off-target populations arise *in vitro*, benefiting the advancement of kidney organoid technologies towards therapeutic outcomes.

## Introduction

The directed differentiation of human pluripotent stem cells (PSC) has been used to form multicellular organoid models of various human tissues^1^, with most protocols following a stepwise differentiation analogous to commitment during mammalian development. While such organoid models represent a major opportunity for accurate human disease modelling and advanced tissue engineering strategies^2,3^, the resulting organoids frequently contain unanticipated, unclassified or ‘off-target’ cell types^4–6^. The temporospatial complexity of embryonic development is such that accurately recapitulating this *in vitro* is likely to result in cellular states that reflect patterning to an adjacent or ‘off-target’ endpoint, the formation of an intermediate cell type not previously characterised or the formation of a completely artificial transcriptional state unique to the *in vitro* culture conditions. Unassigned cell types have been routinely reported in most existing kidney organoid protocols^6,7^. Classification of these cell types remains challenging. While transcriptional atlases of human fetal tissues are rapidly increasing in depth^8,9^, considerable gaps exist. In addition, while many bioinformatic tools exist for cell classification^10–12^, selecting the appropriate reference for comparison with an *in vitro* culture system is also problematic. Even when robust data sets exist, unassigned identities can be observed due to *in vitro* cellular states for which no reference exists.

There are several published protocols for the differentiation of human PSC to kidney organoids containing patterning and segmenting nephrons^7,13–15^. Most kidney organoid protocols are regarded as mimicking gastrulation with commitment to mesoderm via canonical Wnt activation, followed by patterning to IM and induction of nephron formation. Initial patterning can be performed in 2D culture, micro-aggregates or Matrigel^TM^ sandwich cultures (reviewed in Little and Combes, 2019^7^). Consequently, the first week of such protocols is similar to the protocols selected for other mesodermal (heart^16^, blood^17^ and muscle^18^) and neuromesodermal (NMP^19^, neural crest^20^, neuromuscular organoids^21^) cell types. Indeed, the presence of neural, muscle and other mesodermal off-target cell types in kidney organoids has been described previously^4,22–24^, as has considerable variation between batch, cell line and laboratory. While not surprising, the coincident formation of such inappropriate cellular states and their variable presence between differentiations will decrease the endpoint utility with respect to the kidney.

In this study, we classify the cellular identity of all cells present within a comprehensive single-cell RNA-sequencing transcriptional profiling of a commonly applied kidney organoid protocol across 27 days from starting human PSC to kidney organoids. A factorial screen addressing developmental shifts in early mesodermal patterning, the protocol variations included 21 conditions to more comprehensively investigate the source of mispatterning within the canonical protocol. As such, this maximises the potential sources of variation. Prediction of cell identity was achieved with reference to available atlases from mouse and human, including both *in vivo* and *in vitro.* Across the timecourse, 53 individual cellular identities were defined, providing additional clarity to previously unassigned cell types. Using this classification, a reanalysis of all single cell profiling performed using this protocol revealed the presence of the same off-target and indeterminate cellular states, suggesting that these arise irrespective of starting cell or laboratory. Finding and identifying off-target populations within *in vitro* derived tissues has major implications for advancing approaches for future therapeutic applications.

## Results

### Multifactorial timecourse of kidney provides broad transcriptomic landscape for cell identity projection

The screen was designed to capture sample information at the end of each different media composition along the protocol, which we designated as ‘stages’ (Figure 1A). The Takasato protocol was divided into four stages, making single-factor permutations across the first three stages to investigate 21 protocol variations. Stage 1 included 3, 4 or 5 day 7µM CHIR exposures prior to stage 2. At stage 2, we investigated how variations of FGF9, WNT, BMP and RA signalling would impact patterning and organoid development. This included variations of FGF9 at 200ng/mL (F200), FGF9 at 100ng/mL (F), no growth factors (E), or a combination of FGF9 with 2.5µM AGN (FA), 1µM CHIR (FC) or 0.1µM LDN (FL). We additionally investigated the addition of 50ng/ml GDF11 during stage 3 on the control conditions. Technical replicates were pooled together while Hash-tag Oligo (HTO) barcoding was used to multiplex samples for optimal cell sample collection (Figure 1B). Following quality control filtering of the scRNA-seq results, we had collected 150, 957 cells across the entire timecourse, providing the basis of a multifactorial transcriptomic atlas of kidney organoids at four key time-points throughout differentiation (Figure 1C). The whole dataset could be visualized by stage in low dimensions (Figure 1D) which showed minimal overlap between stages. As such, the cell type annotation was performed on each stage independently. Prior to comparing the outcome of each parallel condition, the developing cell types were annotated. While this annotation was more accurate at later organoid stages that have been extensively studied, less-investigated earlier intermediate time points required deeper analyses.

**Figure 1:**
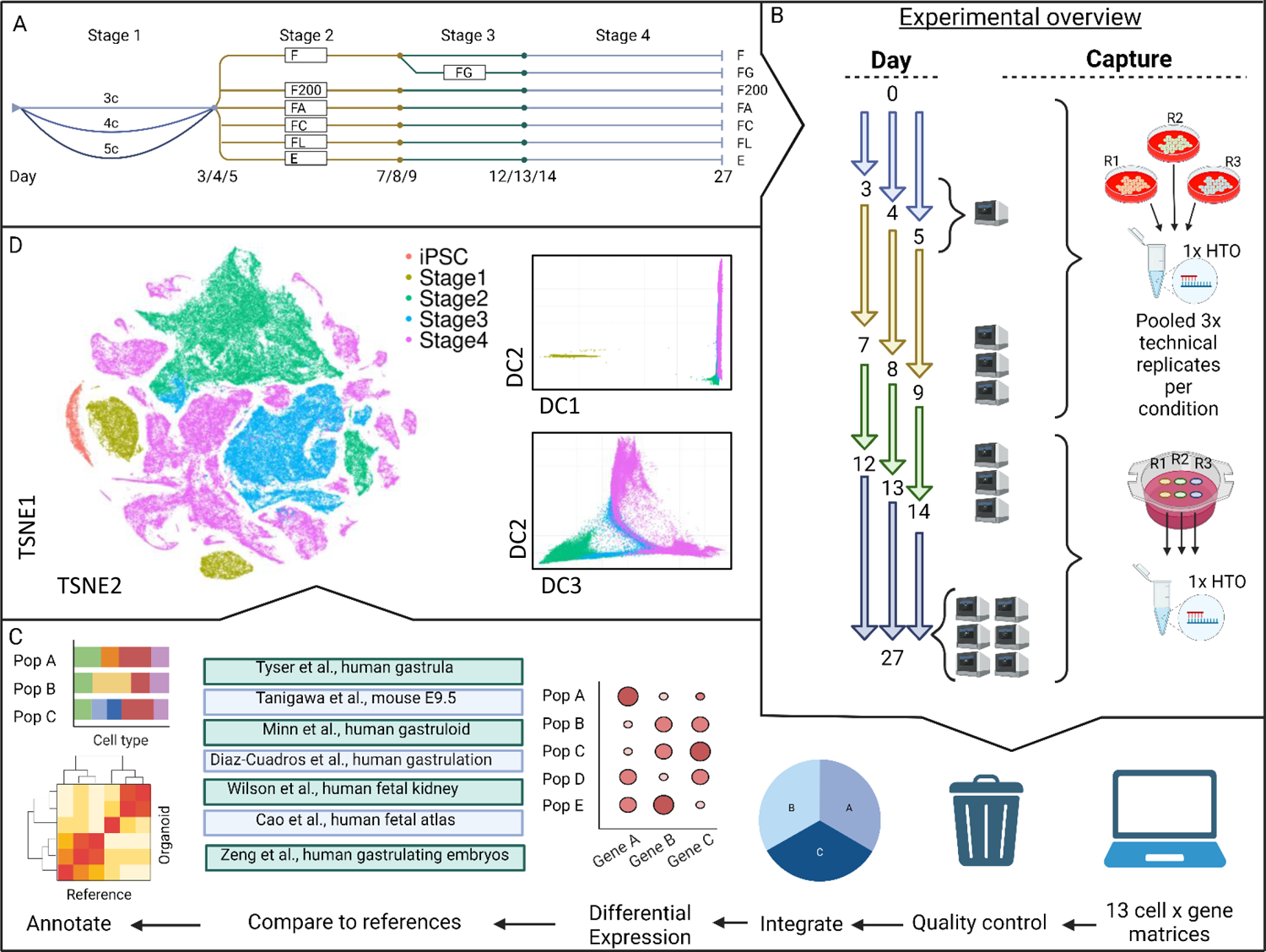
Overview of the experimental outline, execution and data analysis. A) Graphic representation of the kidney organoid protocol used in this study, first developed in Takasato et al.^15^ and then modified in Howden et al.^52^. B) The experimental overview, including the days at which media conditions were changed (numbers at end of coloured arrows), single cell samples were captured (10x machine) and the method of technical replicates and hash-tag oligo labeling. C) The bioinformatics analysis process for cell annotation. D) t-SNE (left) and Diffusion Component (right upper, right lower) representations of the complete organoid timecourse, grouped by protocol stage. The Diffusion Component graphs do not show the iPSC stage. F, FGF9 (100ng/ml); G, GDF11 (50ng/ml); F200, FGF9 (200ng/ml); A, AGN193109 (2.5µM); C, CHIR99021 (1µM); L, LDN193189 (100nM); E, E6; c, CHIR99021 (7µM); t-SNE, t-distributed stochastic neighbour embedding; DC, Diffusion Component; HTO, Hash-tag oligo.

### Reference-based classification of gastrulation-equivalent stages impaired by reference consistency and *in vivo – in vitro* congruence

The process of gastrulation during human development is poorly characterized by single cell transcriptomics, posing a challenge for cell classification within Stage 1. *In vitro*, gastrulation is mimicked via the epithelial to mesenchymal transition from pluripotency to mesoderm. This first stage of differentiation was collected on days 3, 4 and 5 of culture in the presence of CHIR, a GSK3B inhibitor mimicking canonical Wnt signalling and used to initiate mesodermal commitment. Transcriptomic analyses have previously described a primitive streak (PS) identity at the end of Stage 1^15,23^.

We investigated whether reference-based classification with published data sets could identify the most similar cell type for each cell, thereby providing a consensus classification from this. The Tyser^25^ (*in vivo;* human), Tanigawa^26^ (*in vivo;* mouse), Minn^27^ (*in vitro;* human) and Diaz-Cuadros^28^ (*in vtro;* human) reference datasets were used to classify the spanned formation of PS and trunk mesodermal patterning. We utilized label-transfer and machine learning tools (*scPred*^29^*, SCINA*^30^*, SciBet*^31^*, scClassify*^32^*, scAnnotatR*^33^) to classify our data. However, reference-based annotations were highly variable irrespective of the reference dataset or the R-package applied (Figure 2A). Interestingly, batch correction^34^ using Harmony^35^, CSS^36^ or fastMNN^37^ improved but did not remove the batch effect of the human derived single cell references, demonstrating the inherent differences in transcriptional profiles between these reference scRNA-seq data sets (Figure 2A). A Pearson’s correlation using pseudobulk transcriptomes of all references using all DEGs identified (Figure 2Bi) or only the top 25 DEGs from each individual analysis (Figure 2Bii) was still unable to clarify definitive identity.

**Figure 2:**
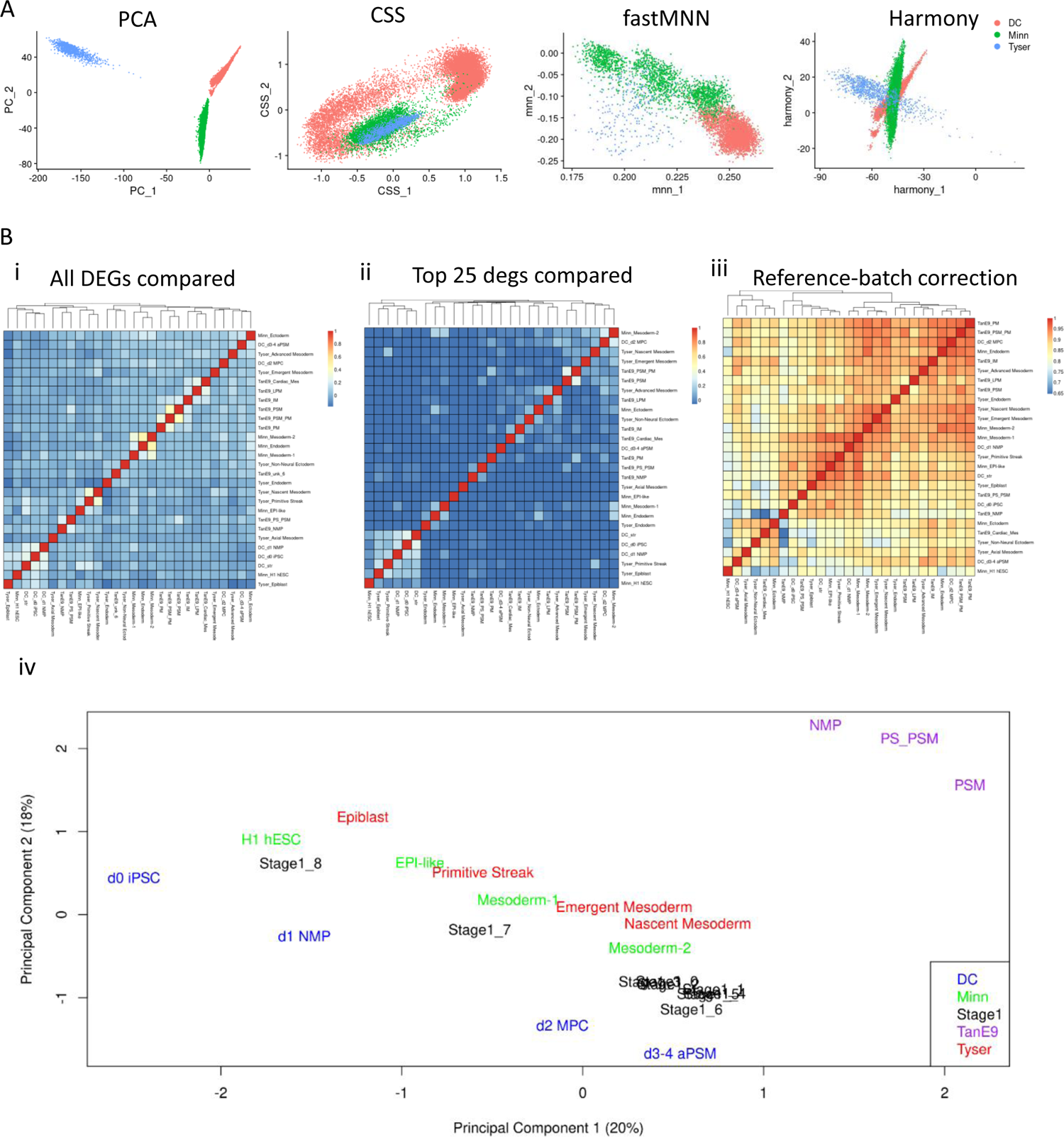
Early embryogenesis human single cell datasets are not optimal for *in vitro* projection. A) 2-dimensional projections of three human gastrulation-stage datasets after dimensionality reduction using PCA, CSS, fastMNN or Harmony to correct for batch effects. No method produces an overlying correlation between the datasets. B) Pearson’s correlation for gene expression between populations within three human gastrulation-stage datasets for i) all DEGs, ii) the top 25 DEGs only, iii) whole transcriptomes after batch correction, iv) PCA projection, this time including clustered stage 1 data from this study.

However, a transcriptomic comparison of all shared genes between references after a batch-correction on the pseudobulked data defined some correlation between references and Stage 1 clusters (Figure 2Biii). The use of disparate nomenclature between studies made the cross-assignment of identity between references difficult. The first two principal components demonstrated similar correlations between the most iPSC-like identities to most differentiated identities across the studies (Figure 2Biv). When applying the models to our data, the results were difficult to interpret. While stage 1 populations were generally regarded as mesodermal lineage, the population identity was disparate and varied (Figure 3A). This repeated for stage 2 populations, where the expected outcomes of nascent or intermediate mesoderm (NM, IM) could be seen if only focusing on one classifier, SciBet (Figure 3B). While this was the most reliable outcome, there was still a large overlap between clusters for identities (Figure 3C). While these outcomes indicated we were indeed generating the expected cell types, lack of clarity in the classifications led us to rely on the standard process of dimensional reduction, clustering and differential gene expression analysis to reliably annotate the cell types present.

**Figure 3:**
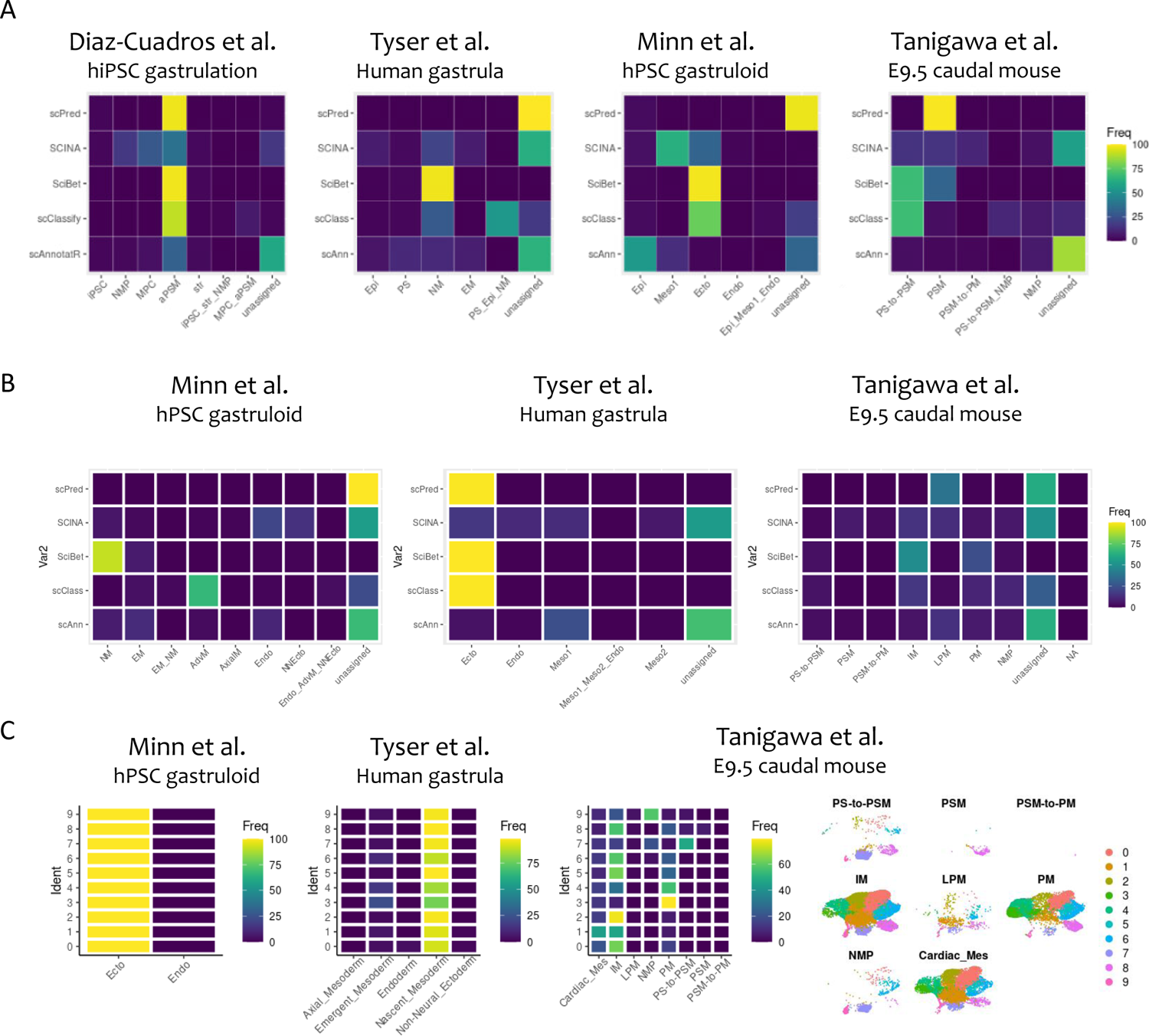
Unbiased cell classification using reference data does not readily resolve populations. A) Heatmap showing the proportion of stage 1 cells classified using a combination of R-based classification algorithms trained on one of four datasets. B) Heatmap showing the proportion of stage 2 cells classified using a combination of R-based classification algorithms trained on one of three datasets. C) Heatmaps and UMAP plot showing the proportion of cells within 10 stage 2 clusters using the R-based classification algorithm SciBet, trained on one of three datasets.

### Incomplete primitive streak differentiation evident during Stage 1

Through projecting cell age onto a UMAP of all stage 1 cells, age-defined groupings were identified (Figure 4A, Table 1) that followed a chronological transition, confirmed using pseudotrajectory analysis (Figure 4B). The two smallest clusters expressed pluripotent markers *EPCAM, POU5F1* and *SOX2*, with one cluster also expressing differentiation markers *SP5, MIXL1, FGF8* and *FGF17* (Figure 4C), indicating iPSC-like and neuromesodermal progenitors (NMP) respectively. The larger remaining clusters expressed PS markers alone and could be divided into three groups, giving five populations at this stage (Figure 4C). The iPSC and NMP clusters contained cells of all three ages, indicating these populations persisted regardless of extended CHIR exposure (Figure 4D). Pseudotime projection confirmed cells were transitioning from the 3c population to 5c (Figure 4E). Distinguishing between them was the increasing expression of *MIXL1, MSX2* and *TBX6* with age (Figure 4C,F). As these populations all expressed the key markers for posterior primitive streak (pPS) they were distinguished as early_pPS, late_pPS_1 and late_pPS_2 based on their position along the correlating pseudotime-age axis.

**Figure 4:**
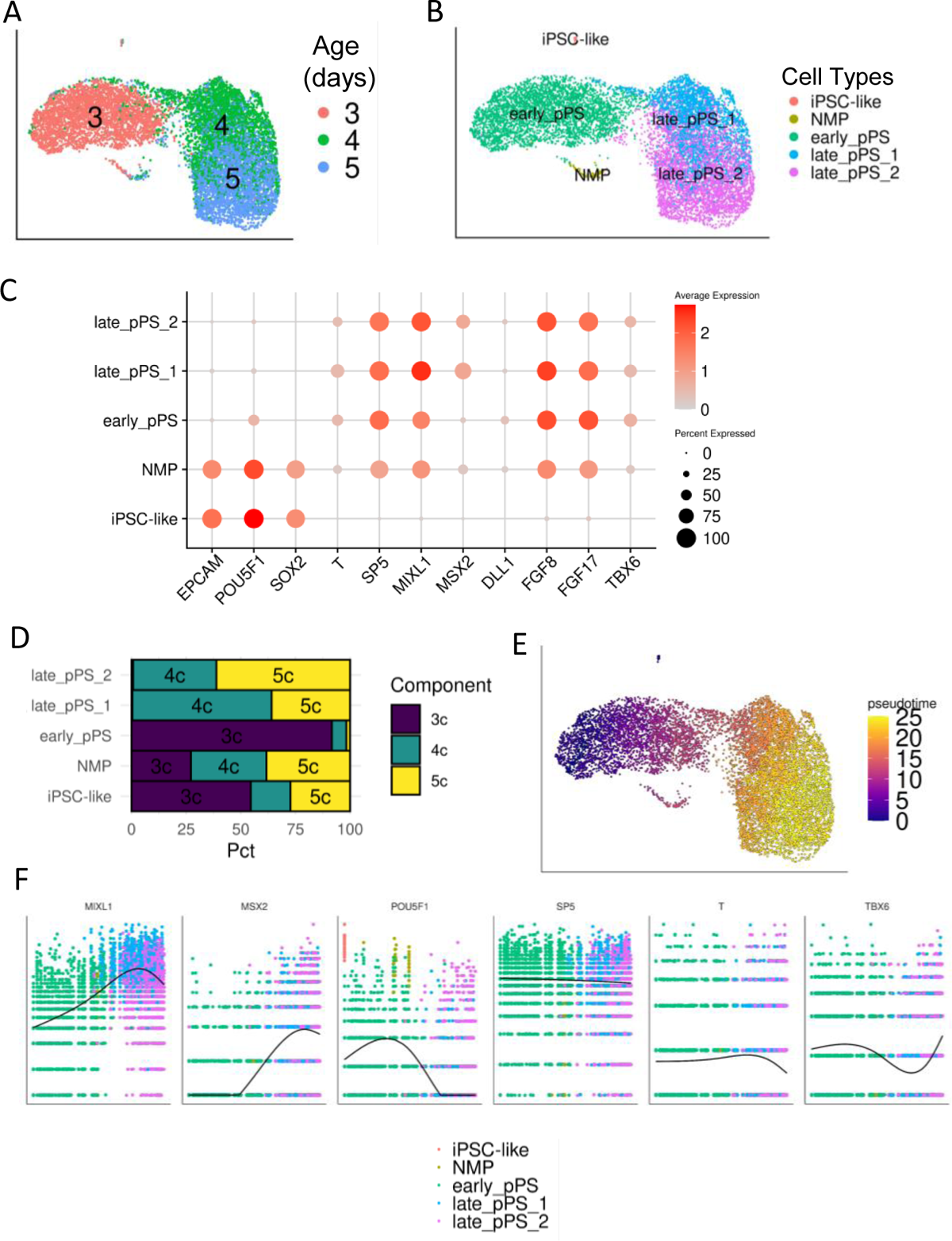
Cell transition to primitive streak using CHIR includes asynchronous differentiation and lagging iPSC and NMP cells A) UMAP representation of Stage 1 cells grouped by the sample age in days. B) UMAP representation grouped and labelled by annotation of stage 1 cells into iPSC-like, NMP, early_pPS, late_pPS_1, late_pPS_2. C) DotPlot showing the expression of known markers of pluripotency, neuromesodermal progenitors, primitive streak and early presomitic mesoderm. Dot size relates to proportion of cells in cluster expressing the gene, colour scale shows the relative level of expression. D) Proportion of cells from each day of differentiation within each cell type. E) UMAP representation of the pseudotime across stage 1 differentation. F) Change in expression of key markers of differentiation across time during stage 1 differentiation. UMAP, uniform manifold approximation and projection; PS, primitive streak; NMP, neuromesodermal progenitors; PSM, presomitic mesoderm; NC, neural crest; NP, neural progenitors; M, mesoderm; IM, intermediate mesoderm; PM, paraxial mesoderm.

**Table 1.** Parameters used for dimensional reduction.

| Stage | Genes | Batch correction | Components used for UMAP / TSNE | Cluster resolution |
| --- | --- | --- | --- | --- |
| Stage 1 | 3000 | NA | 30 PCs | 0.5 |
| Stage 2 | 3000 | NA | 30 PCs | 0.3 |
| Stage 3 | 3000 | NA | 30 PCs | 0.5 |
| Stage 4 | 3000 | NA | 15 PCs | 0.3 |
| All | 3000 | CSS (default settings) | 40 PCs / 50 PCs | NA |

The presence of cells that had not fully committed to posterior primitive streak indicated that there had been incomplete differentiation from the CHIR addition. The proportion of cells remaining in an iPSC-like state decreased with time, but did not deplete by 5 days of treatment. In contrast, early_pPS consisted of >95% 3 day cells, while late_pPS_2 had >50% 5 day cells and almost no 3 day cells. This demonstrated that a population of cells are either unable to differentiate in response to CHIR, or there is an asynchronous differentiation that is identifiable at 5 days of CHIR-mediated differentiation.

### Stage 2 contains a heterogeneous mix of mesodermal and neural cell identities

Patterning from PS to intermediate mesoderm (IM) was hypothesized to be the most critical window for influencing endpoint cell types. In the original Takasato protocol, this is performed in 2D culture with IM induced via the addition of FGF9^15,38^ (F200) (Figure 1A). The combined Stage 2 dataset contained 44,282 cells at 7, 8 or 9 days of culture. There was some disparity in cell distribution when comparing the samples from 3, 4 or 5 days of CHIR exposure (Figure 5A). As there was no consensus found when applying automated classifiers as described above (Figure 3), dimensional reduction, clustering and differential gene expression analyses were employed to identify the cell types present.

**Figure 5:**
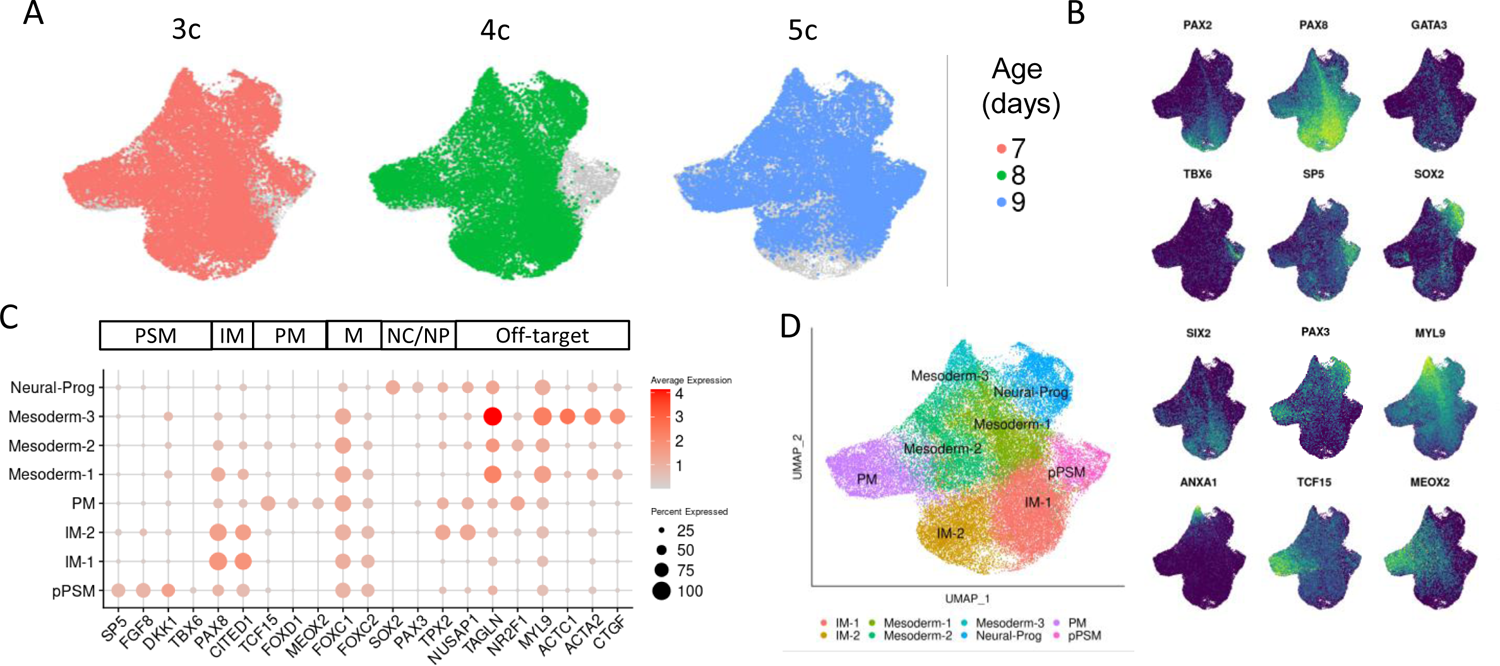
Heterogeneous cell populations present after stage 2 of differentiation indicate off-target populations can be identified early in organoid development A) UMAP representation of stage 2 cells coloured by the length of initial Wnt agonism during stage 1 patterning, with unincluded cells coloured grey. B) Expression of population marker genes plotted onto the UMAP representation of stage 2 cells. C) DotPlot showing expression of key markers of PSM, IM, M, NC/NP and off-targets in each annotated population. Dot size relates to proportion of cells in each cluster, colour scale shows the relative expression level. D) UMAP representation of Stage 2 data indicating final cell identity annotations. UMAP, uniform manifold approximation and projection; PS, primitive streak; NMP, neuromesodermal progenitors; PSM, presomitic mesoderm; NC, neural crest; NP, neural progenitors; M, mesoderm; IM, intermediate mesoderm; PM, paraxial mesoderm.

Kidney-deriving IM populations are known to express *PAX8*, *PAX2* and *GATA3*, while nephron progenitor cells themselves (NPCs) are defined by *SIX2* expression^39^. Plotting the expression distribution of these and other markers of potential cell populations shows the heterogeneity of cell types present at this stage (Figure 5B). The presence of paraxial mesoderm (PM) markers *TCF15* and *MEOX2* indicated a shift in mesoderm patterning in one or more conditions. The overlapping expression of *PAX8* and *SIX2* indicated there were definitive kidney lineage cells present. However, the localised expression of neural (*SOX2*) and muscle (*MYL9*, *ANXA1*) precursors indicated off-target populations could be identified at this time point.

Comparing the top DEGs in each cluster (Figure 5C) indicated patterning to *PAX8^+^CITED1^+^* IM (IM-1 and IM-2), *TCF15^+^MEOX2^+^*PM, *FGF8^+^SP5^+^*posterior presomitic mesoderm (pPSM) and *SOX2^+^ PAX3^+^* neural progenitor or neural crest (Neural-Prog). All clusters except Neural-Prog were enriched for *FOXC1* and *FOXC2* indicating a commitment to mesoderm^40,41^ (Figure 5C). Three populations that did not have more specific lineage or population markers were labelled Mesoderm-1,2 and 3 (Figure 5D).

### Optimisation of DevKidCC to accurately predict kidney and non-kidney cell types in complex *in vitro* datasets

After the 2D culture to Stage 2, all cells were dissociated and reaggregated to form a 3D micromass and cultured at an air-media interface (Transwell^TM^ culture)^15^. In the subsequent 5 days of culture, during which FGF9 remains present, NPCs should arise and begin transitioning to early nephron (EN) via a mesenchymal-to-epithelial transition^24^. Classification of this stage, containing distinguishable kidney cell types, was initially performed using *DevKidCC*^6^, a classification tool containing a hierarchal set of binary models trained on a multi-sample human fetal kidney (HFK) reference. Based on s*cPred*^29^, the tool *DevKidCC*^6^ uses three classification tiers to provide a definitive identity to a cellular state in a human fetal kidney.

Initial application of *DevKidCC* showed that a substantial component of cells within published kidney organoid datasets did not match cells present *in vivo* (classified as unassigned) (Figure 6A). Of the 34,386 cells collected at stage 3, 52.1% were classified with a kidney lineage while 47.9% remained unassigned. The 52.1% of kidney cells subset into 31.5% nephron lineages, while 20.6% were mesenchymal lineages (Figure 6A). No cells were defined as endothelial while five cells of ureteric epithelial identity were classified. Plotting each lineage separately displayed distinct groupings of nephron, NPC and stroma (Figure 6A). However, large overlapping populations of NPC-like and unassigned cells were also present. *DevKidCC*^6^ had previously assigned the term NPC-like to cells with a transcriptional congruence similar to NPC, but below the default threshold of transcriptional congruence (set at 0.7) and lacking expression of *PAX2*^6^. Cells identified as NPC-like within this stage displayed a broad distribution across stroma and unassigned domains suggesting that this classification represented an artefact (Figure 6A). The large proportion of unassigned cells at Stage 3 also suggested that, during this transition from IM to nephron, this classification tool was suboptimal. The expression of known nephron and non-nephron markers indicated that, while the NPC and nephron lineages had been accurately identified, there were false negatives in the unassigned cells. Further, the population of *SOX2* expressing off-targets had been classified heterogeneously as nephron, NPC-like or unassigned (Figure 6B).

**Figure 6:**
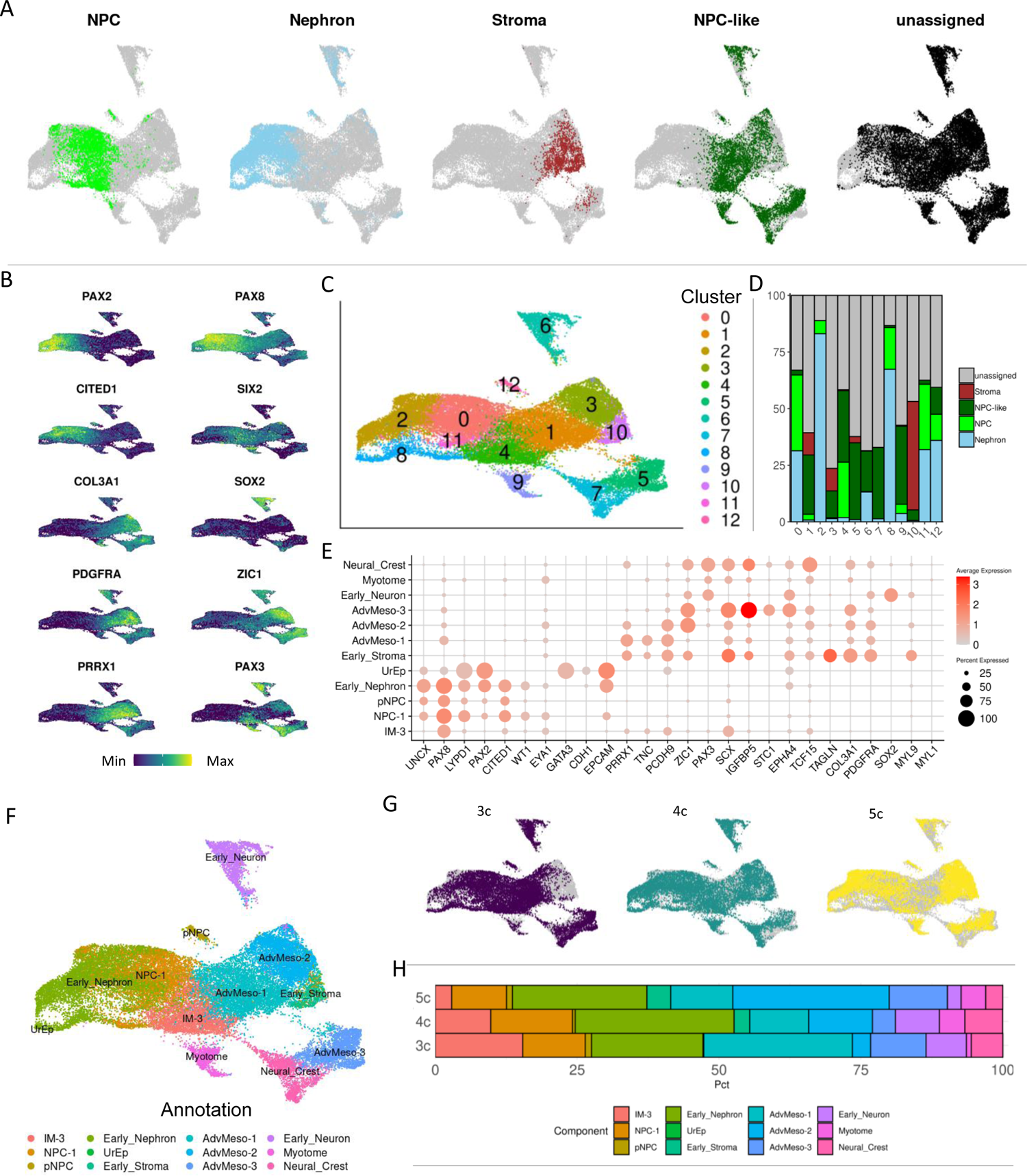
Kidney and ‘off-target’ cell types are present at Stage 3 of differentiation. A) UMAP representation of the Stage 3 cells grouped and split by *DevKidCC* classified identity, with all unincluded cells in grey. B) UMAP representation of Stage 3 cells coloured by gene expression of *PAX2*, *PAX8*, *CITED1*, *SIX2, COL3A1, SOX2, PDGFRA*, *ZIC1, PAX3* and *PRRX1*. C) UMAP representation of Stage 3 cells grouped by cluster identity. D) Bar chart showing the proportion of *DevKidCC* classified lineage identities in each of the 13 clusters. E) DotPlot showing the expression of key marker genes in Stage 3 clusters. Dot size relates to proportion of cells in each cluster, colour scale shows the relative expression level. F) UMAP representation of Stage 3 cells grouped by annotated identity. G) UMAP representation of Stage 3 cells grouped and split by length of CHIR during stage 1. H) Bar chart showing the proportion of annotated populations present within each length of CHIR during stage 1. UMAP, uniform manifold approximation and projection; PM, paraxial mesoderm; NMP, neuromesodermal progenitors; IM, intermediate mesoderm; MM, metanephric mesenchyme; EN, early nephron; EDT, early distal tubule; EPT, early proximal tubule; PEC, parietal epithelial cells; EPod, early podocytes; Pod, podocytes; Meso, mesoderm; tNC, trunk neural crest; NP, neural progenitors.

To readdress this, the Stage 3 data was clustered into 12 populations using k-means (Figure 6C) with each cluster subsequently reclassified using *DevKidCC* (Figure 6D). In many clusters, unassigned and NPC-like cells were the most prevalent identity. Within clusters 0, 2, 8, 11 and 12, most cells were classified as nephron or NPC identity. These clusters expressed IM and metanephric mesenchyme (MM) markers *CITED1*, *PAX2* and *PAX8,* whereas the remaining clusters expressed non-nephron markers such as *PDGFRA*, *ZIC1* and *PRRX1* (Figure 6E).

Interestingly, *PAX2* seemed to be more specific to the nephron lineage, while *PAX8* was expressed across a broader range of cells, including coexpressed with *ZIC1, SOX2* and *PAX3* populations (Figure 6B,E).

Cluster 6 was enriched for the neural markers *SOX2* and *EPHA4,* indicating these were a neural off-target population. Cluster 9 expressed *MYL1* strongly in a few cells, alongside *TCF15*, *PAX3* and *SCX* indicating a muscle-progenitor population we describe as myotome (Figure 6E).

Clusters 1, 3 and 10 displayed a similar DEG expression profile and included cells designated as stroma by *DevKidCC* (Figure 6A,F). Mesenchymal genes *PDGFRA* and *COL3A1* were expressed, as was the muscle lineage determinant *PRRX1*, but no other indicative markers of a kidney specific population (Figure 6E). These were annotated as early_stroma and advanced mesoderm (AdvMeso-1/2).

A mix of gene expression is present in clusters 5 and 7. Weak expression of *PAX8, SIX2* and *ZIC1* is present in cluster 5, with *PRRX1, SOX2* and *PAX3* in cluster 7 (Figure 6E). This profile predicts that these cell populations are similar to mesoderm and neural crest respectively.

These analyses suggested that lineage divergence between nephron and non-nephron occurs during stage 3 of differentiation and enabled cell identities to be assigned to all clusters (Figure 6F). These included ‘on-target’ cell types (Nephron, NPC, Stroma), a proliferating nephron cell type (pNPC), unspecified or ambiguous mesenchymal populations (IM-3, AdvMes-1/2/3), and likely ‘off-target’ populations (Neural crest, early_Neuron, Myotome). Contribution to all clusters was seen irrespective of initial CHIR duration (Figure 6G), although 3c contained fewer ‘on-target’ stromal cells. There were differences in the prevalence of non-kidney cell types depending upon the duration of initial CHIR patterning, with AdvMeso-1 being enriched in 3c while AdvMeso-3 was enriched in 5c (Figure 6H). Only early nephron populations around the stage of the comma-shaped body, an early nephron structure, were identified.

### Development of neural and muscle ‘off-targets’ in later organoids

At Stage 4 (day 27 organoids), 59,631 cells were collected for single-cell profiling. As this stage resembles a trimester 2 human fetal kidney^15^, it is optimal for *DevKidCC* classification. The nephron lineages (NPC, Nephron, UrEp) were clearly distinguished from the mesenchymal (Stroma, NPC-like) and unassigned cell types (Figure 7A). A single endothelial cell (Endo, not shown) and few UrEp cells were present, which agrees with previous studies^4,6,23,42–44^. We repeated the process of k-means clustering and analysis of *DevKidCC* per cluster, for more accurate classification (Figure 7B). This method identified nephron (clusters 3, 5, 6, 7, 13), stromal (clusters 0, 9, 12, 14) and unassigned (clusters 1, 2, 4, 8, 10, 11) cell types (Figure 7C).

**Figure 7:**
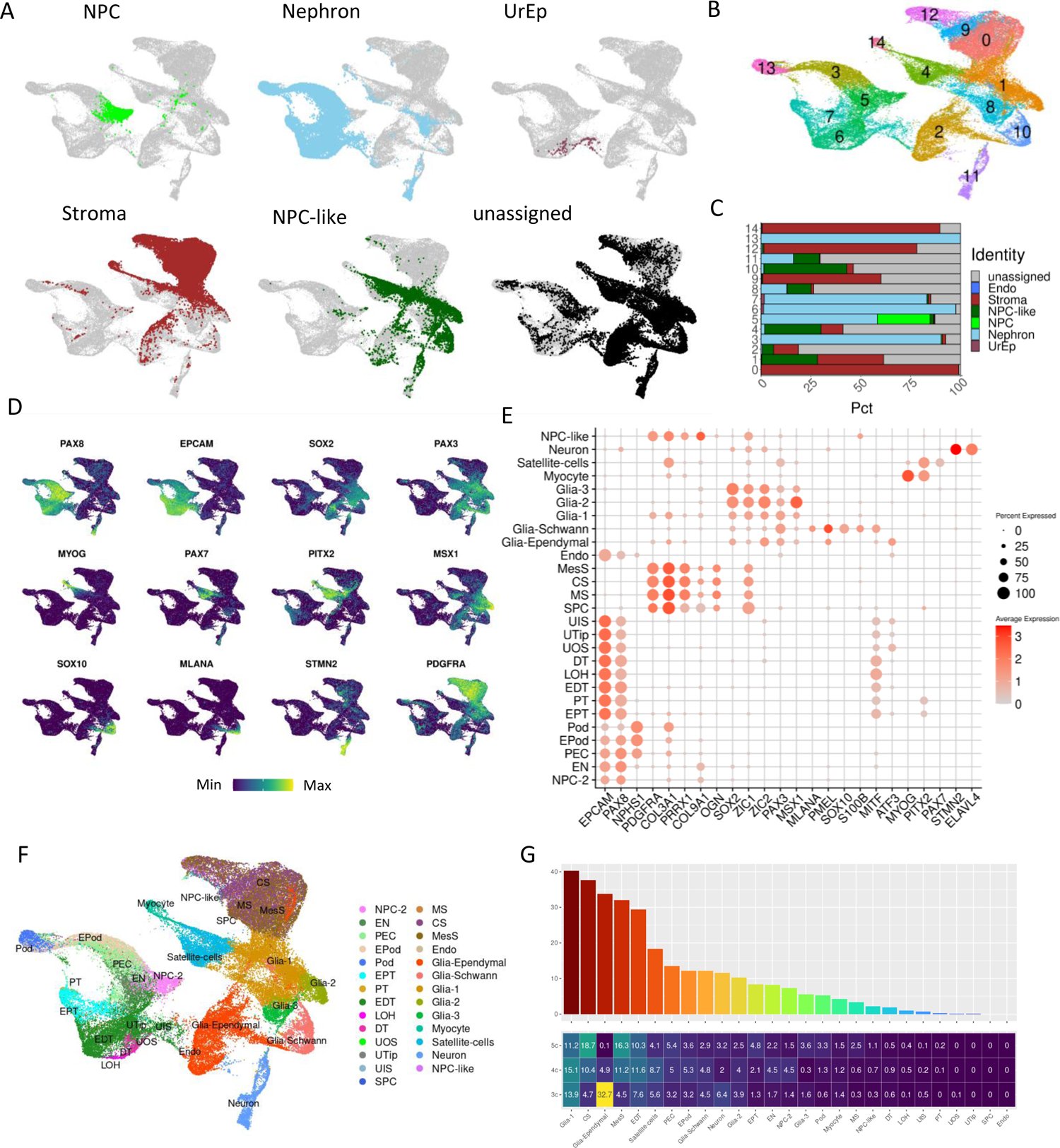
Kidney and ‘off-target’ cell types present in final kidney organoids (Stage 4). A) UMAP representation of Stage 4 cells grouped and split by lineage identification using *DevKidCC*. B) UMAP representation of Stage 4 cells clustered into 15 populations. C) Bar chart showing the proportion of *DevKidCC* lineage classification grouped by cluster identity. D) UMAP representation of Stage 4 cells showing the expression of markers that are regionally enriched in nephron and non-nephron populations. E) DotPlot showing expression of key population markers used in classification. Dot size represents proportion of cells in cluster expressing gene, colour represents average expression level. F) UMAP representation of Stage 4 annotated cell types. G) Heatmap and bar chart combination to show the proportion of each population contribution to the organoids from the 3c, 4c and 5c conditions (all growth factor combinations combined) H) UMAP (left) and heatmap/bar chart combination (right) showing the samples from the original takasato protocol only. UMAP, uniform manifold approximation and projection; DEGs, differentially expressed genes; NPC, nephron progenitor cells; UrEp, ureteric epithelium; Endo, endothelium; UTip, ureteric tip; UOS, ureteric outer stalk; UIS, ureteric inner stalk; PT, proximal tubule; Pod, podocytes; PEC, parietal epithelial cell; LOH, loop of Henle; EPT, early proximal tubule; EPod, early podocyte; EN, early nephron; EDT, early distal tubule; DT, distal tubule; SPC, stromal progenitor cells; MS, medullary stroma; MesS, mesangial stroma; CS, cortical stroma.

*PAX8* and *EPCAM* were definitively expressed in the nephron populations, although weakly in the podocytes as expected (Figure 7D,E,F). The top DEGs in the unassigned clusters related to neural (*SOX2, PAX3, STMN2)*, neural crest/pigment (*MLANA, SOX10, MSX1)*, muscle (*MYOG, PAX7, PITX2)* and general mesenchymal (*PDGFRA)* genes (Figure 7D). Much of the stromal population co-expressed mesenchymal genes, *PDGFRA*, *COL3A1* and *COL9A1,* and muscle lineage marker, *PRRX1,* in addition to chondrocyte lineage marker, *OGN* (Figure 7E). A number of populations expressed variations in neural and glial markers. In particular, one population expressed *ATF3*, a marker of ependymal cells (Figure 7E), not previously identified in kidney organoid studies. Interestingly, the ependymal cells were not present in the 5 day CHIR exposure conditions (Figure 7F). The pigment-expressing cells (*MLANA*, *PMEL*) are most likely Schwann cells as they also express neural crest and glial markers (*S100B*, *SOX10*). These have previously been labelled as melanocytes in other studies^4,5^. Additionally, 4c and 5c conditions contained more cells classified as kidney lineages (Figure 7G).

Given the large proportion of non-kidney cell types present at this stage, label transfer using the tool Azimuth^47^ was utilised to compare all cells at stage 4 with a human fetal atlas containing cell types from 15 organs between 72 to 129 days post-conception^9^. When investigating the distribution of maximum similarity scores at stage 4, all on-target clusterers cells showed highest correlation with a Kidney-Metanephric and Kidney-Ureteric bud classification (Figure 8A).

**Figure 8:**
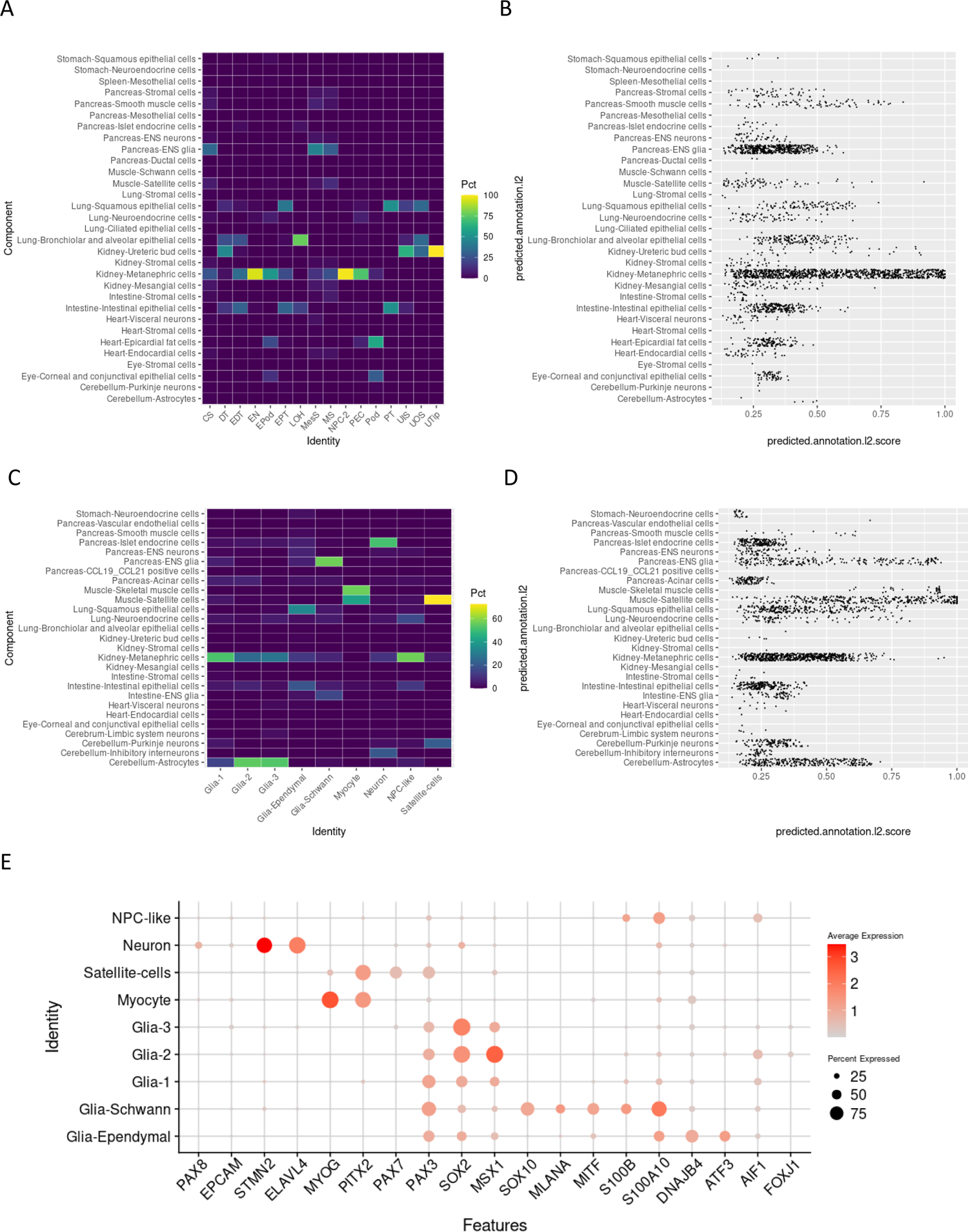
Comparison of on-target and off-target populations within Stage 4 with human developmental atlas using Azimuth. A) Heatmap showing the proportion of cells in each Stage 4 cluster for all *on-target* cells (CS, DT, EDT, EN, EPod, EPT, LOH, MesS, MS, NPC-2, PEC, Pod, PT, UIS, UOS, UTip) in Stage 4 when compared to the human developmental atlas^9^ using the Azimuth^47^ web browser. B) Scatterplot showing the distribution of maximal predicted scores per cell type in A. C) Heatmap showing the proportion of cells in each Stage 4 cluster for all *off-target* cells (Glia-1, Glia-2, Glia-3, Glia-Ependymal, Glia-Schwann, Myocyte, Neuron, NPC-like, Satellite-cells) in Stage 4 when compared to the human developmental atlas^9^ using the Azimuth^47^ web browser.. D) Scatterplot showing the distribution of maximal predicted scores per cell type in C. E) DotPlot showing the expression of off-target markers correlating with off-target populations. Dot size relates to the proportion of cells in each cluster, colour scale shows the relative expression level.

However, for most cells the predicted score was <0.5 (Figure 8B) indicating only a weak similarity to any population within the reference. It was interesting to see that stromal populations had very little correlation to the Kidney-Stromal reference, while podocytes (Pod) no correlation with kidney identity within this Azimuth dataset (Figure 8A,B). When comparing the off-target populations to the reference, many cells retained a strongest correlation to Kidney-Metanephric although most cells had less than 0.5 mapping score (Figure 8C,D). The gene expression of key markers from the off-target populations correlate with the label transfer, providing confirmation of these *in vitro* cell identities (Figure 8E).

These data demonstrated the large proportions and varieties of off-target cell types in kidney organoids. Further, the comparison between marker gene and label transfer analyses highlight the difficulty in reference-based identity assignment of *in vitro* data onto *in vivo* references.

### Off-target populations vary in data from previous studies

Determining the extent to which these off-target populations arose from the variations included in the factorial screen required comparison back to the initial protocol (F200) (Figure 9A). In organoids generated using the original Takasato protocol (F200), there was an increase in off-target neural populations in the 3c condition, while the 4c and 5c contain a much better contribution of on-target stroma and nephrons (Figure 9A). As such, the control organoids in this study are comparable to previous datasets. Further, the atlas as a whole provides valuable information for how growth factor modifications can impact organoid development.

**Figure 9:**
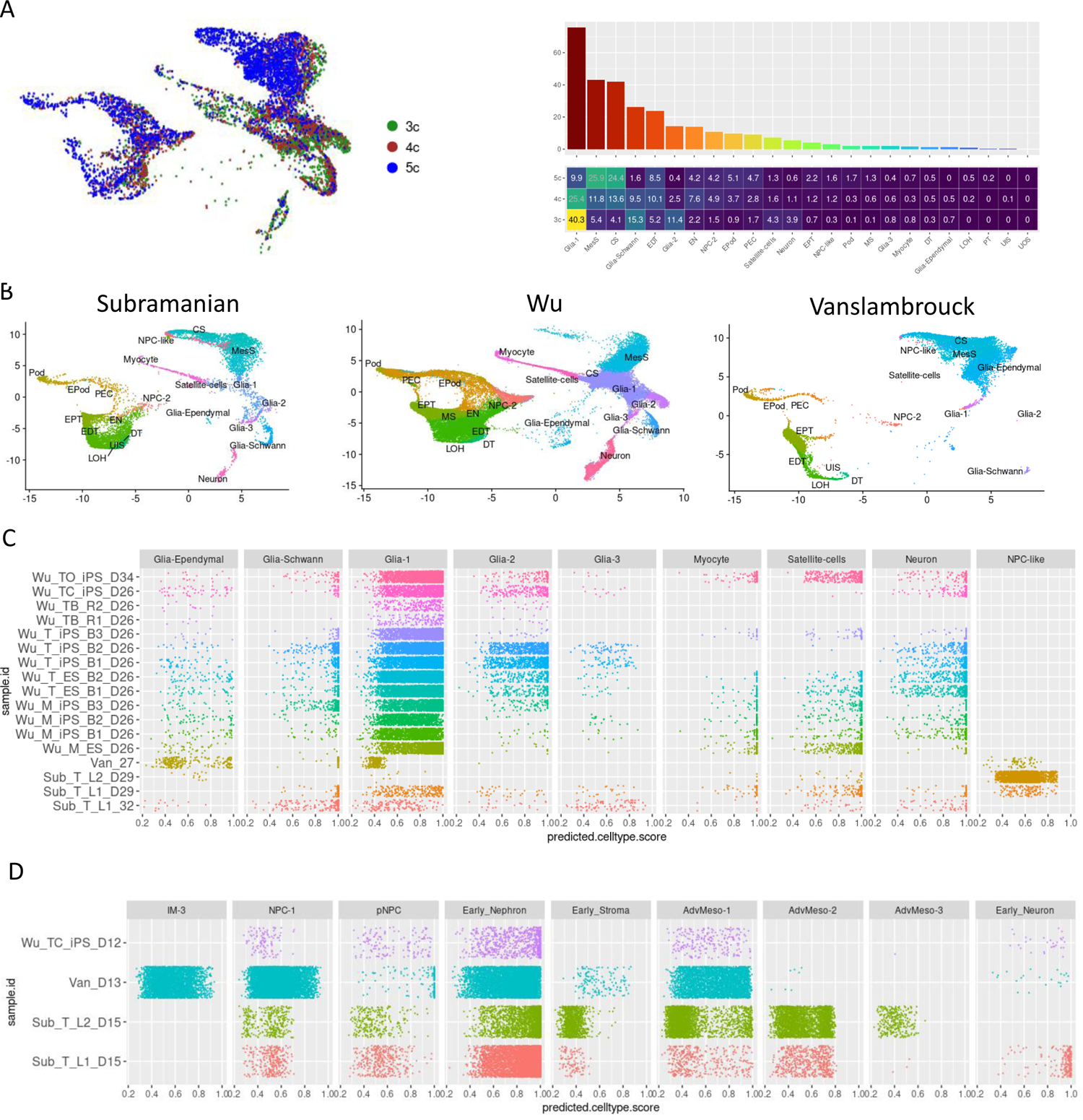
Organoid cell proportions within previous studies when projected onto this study’s atlas show consistent off-target populations. A) UMAP representations of previous studies showing the distribution of cell types in comparison to the stage 4 data in this study. B) The maximum projected similarity for all cells in each sample from the three previous studies at timepoints matching stage 4. C) The maximum projected similarity for all cells in each sample from the three previous studies at timepoints close to stage 3.

The observation of off-target cell types within the Takasato^15^ protocol has been reported previously^4–6^. A direct comparison was made between the transcriptional profile of classified off-target cells in this study with published data from two prior studies using the same protocol (Subramanian et al, Wu et al)^4,5^ and a more recent modification which reports enhanced proximal tubule maturation (Vanslambrouck et al)^48^. These published protocols used distinct cell lines but collected data at similar timepoints to Stage 3 and Stage 4. Based on the prior observation of neural off-targets, the Wu dataset also contained transcriptional data from cultures treated with an inhibitor of BDNF to reduce neural commitment. Using Seurat, datasets from each publication were labelled based on the highest matched cell type using the anchor label transfer method. The most mature organoids from each study (compared to Stage 4) showed distinct differences in the distribution and proportion of off-target markers. Subramanian^5^ organoids contained more distalised nephrons and fewer neural subsets (Figure 9B). All lines, batches and treatments included in the Wu^4^ dataset also contained the cell types described in our study (Figure 9B). Conversely, the data from Vanslambrouck^48^ lacked almost all neural and muscle off-targets, although was previously shown to contain a cartilage population not detected using the atlas in the current study (Figure 9B). The distribution of maximum probability scores for all off-target cells in these samples, separated by individual conditions per study, confirmed that most cells were above 0.5 (50%) score for similarity to their assigned identity (Figure 9C). The decrease in neural cell types, both in the Vanslambrouck (Van_D27) and Wu inhibition of BDNF (Wu_TB_R1_D26 and Wu_TB_R2_D26) impacted all neural subtypes (Figure 9C). Less prevalent off-target populations in our dataset (Ependymal, Schwann, neurons and myocytes) were identified in some but not all conditions (Figure 9C).

This analysis was repeated for the stage 3 samples in each of these studies (Figure 9D). At stage 3, there were predictable differences in cell projections between samples based on the protocol used in the study. The Vanslambrouck day 13 sample was maintained in a nephron progenitor expansion media, which is reflected in the increase in higher scores for IM-3 and NPC-1 at this time. In contrast, the other studies had initiated nephrogenesis and had higher proportions of differentiated cell types, including the ambiguous mesoderms AdvMeso-2 (Figure 9D).

The consistent generation of off-target cell types shows that a spectrum of cellular heterogeneity is inherent in the differentiation protocol. These can be manipulated via targeted changes to optimise nephron maturation^48^ or inhibit unwanted genetic programs^4^. As such, this study provides a valuable database that can be used to advance kidney organoid differentiation.

## Discussion

In humans, the permanent kidney arises from posterior IM, with this in turn arising from the posterior primitive streak. In the mammalian embryo, the IM gives rise to three pairs of nephron-containing organs, the pronephros, mesonephros and metanephros. More anterior IM, regarded as exiting the tailbud at an earlier timepoint, gives rise to the pronephros, mesonephros and nephric duct while posterior IM forms the metanephros. Across the mediolateral axis of the mammalian embryo, the mesoderm is patterned from paraxial through intermediate to lateral plate mesoderm. The PM forms from an NMP population co-expressing *SOX2* and the mesodermal marker *Brachyury*/*T*. NMP can give rise to neural cell types of the spinal cord or PM, which gives rise to somites. All these cellular states arise from gastrulation. Hence, to accurately recreate the developing metanephros from pluripotent stem cells requires exquisite mesodermal patterning.

In this study, a complete temporal factorial screen based around the most commonly used kidney organoid protocol was thoroughly interrogated in order to classify the cellular identity of all cells present across time and condition. This analysis has provided a thorough evaluation of the ‘on-‘ and ‘off-‘ target populations present within kidney organoids generated using the Takasato protocol and variations within this study. We identified nephrons at both immature (NPC-2, EN, EDT, EPT, EPod) and mature (DT, LOH, PT, PEC, Pod) stages, some evidence of ureteric lineages (UTip, UIS, UOS) in the shorter CHIR variants and populations of kidney specific stroma (CS, MS, MesS). A thorough investigation of the NPC-like and unassigned cells within the organoids identified populations of neural (Neuron, Glial subtypes) and muscle (Myocyte and Satellite-cells) while some undecided cells do retain a NPC-like identity.

scRNA-seq is often analysed by reducing the dimensional information from thousands of cells and genes into tens of component dimensions in an attempt to remove noise and hone in on the patterns of gene expression that explain biological context^49^. Cells are then grouped using clustering algorithms based on these lower dimensions prior to visualization, often after further dimensional reduction^50,51^. While appropriate for the definitive assignation of identity to mature distinct populations this is problematic with developing tissue and more so with an *in vitro* model of a developing tissue. Hence, the central challenge in this study was the accurate assignment of cellular identity in the absence of a ground truth. This was a problem both early, where data on gastrulation and early mesoderm patterning was limited, and late, with the presence of ‘off-target’ populations with no clear *in vivo* counterparts. We found that trying to directly compare whole or even partial transcriptomes to the few available reference datasets for Stages 1 and 2 was difficult due to the variability in the transcriptional profiles reported and the nomenclature used in each study. Regardless of the method used for reference-based classification, the results were unreliable. It proved more effective to rely on selected expressed genes to provide the best fit to a reference population. This illustrated that there is room for improvement in reference-based classification of early development.

At later stages, initial classification deployed *DevKidCC*^6^. This tool is based on an organised set of binary models trained on *in vivo* human fetal kidney data to provide a robust and unbiased prediction kidney cellular identity, enabling direct cross comparison between datasets^6^. When applied to Stages 3 and 4, the classification of nephrons was highly accurate, but there was increased margin of error with stroma and unassigned (off-target) populations. Our analysis also suggested that the classification of NPC-like was an *in vitro* artefact and should be reconsidered. Classification based on DEG, Pearson’s correlation and label transfer using Azimuth with a general human developmental reference enabled a more comprehensive assignation of likely cell identity for all non-nephron cells at these stages. Again, it should be noted that the sparsity of scRNA-seq data available, use of arbitrary clustering and the presence of likely *in vitro* transitional cell states will remain a challenge to definitive cell classification.

This study represents a comprehensive temporal single-cell transcriptional atlas across *in vitro* differentiation from pluripotency through mesoderm patterning to kidney. The purpose of this manuscript was to provide a classification of cellular identity for all cells present within an *in vitro* differentiation to human kidney. What was observed were distinct ‘off-target’ cell states present alongside on-target kidney cell types. The classification of all 150,957 cells into 53 distinct identities. The resulting atlas will serve as a valuable resource for the kidney organoid community, enabling a thorough dissection of where and why current kidney organoid protocols are imperfect and serving as a reference upon which modifications of the protocol can be compared.

## Acknowledgements

This work was supported by the Australian Research Council (SR1101002; DP190101705), the National Institute of Health (DK107344) and the National Health and Medical Research Council (GNT1136085). JEP is supported by the National Health and Medical Research Council Fellowship (APP1107599) and with the support of the Fok Family in memory of Dr. and Mrs. Wing Kan Fok. MHL and SBW are supported by the Novo Nordisk Foundation Center for Stem Cell Medicine (<u>NNF21CC0073729</u>). We acknowledge MCRI Operational Infrastructure Support and the Stafford Fox Medical Research Foundation MCRI Genome Editing Facility for generating pluripotent stem cell lines. We thank the Garvan Centre for Cellular Genomics for assistance with experimental planning, 10x single-cell library preparation, sequencing and data curation.

## Author contributions

SBW, MHL and JEP conceived the study. SBW, JMV, AM and SEH performed kidney organoid differentiations. SBW, AA and DN performed bioinformatic analysis with guidance from JEP and AT. SBW and MHL wrote the manuscript while all authors assisted in manuscript preparation. All authors read and approved the final manuscript.

## Methods

### Kidney organoid differentiation

The iPSC line used in this study was CRL2429-MAFB^mTagBFP2^/GATA3^mCherry^ ^53^, maintained and expanded at 37°C in 5% CO_2_ in Essential 8 medium (Thermo Fisher) on Matrigel^TM^-coated (Thermo Fisher) plates with daily media changes and passaged every 3-4 days with EDTA in 1x PBS buffer as described previously^54^. An initial seeding density of 7 x 10^4^ cells into 6-well culture plates was then cultured with variations based on the Takasato protocol^15^ modified to use TeSR-E6 medium^52^ (STEMCELL technologies) further described in the partner manuscript (Wilson et al.,). Briefly, cells were cultured in 1.5mL of TeSR-E6 with 7µM CHIR99021 (Tocris) for 3-5 days. Then cells were cultured in 2mL of TeSR-E6 with one of six growth factor combinations for 2-4 days. Cells were then dissociated and resuspended into a dense 200k/µl cell paste with TeSR-E6. Each organoid was generated by transferring 1µl onto the top, air-facing side of a Transwell^TM^ membrane (Corning), with six organoids per Transwell. A pulse of WNT signalling was provided using 1.1mL of 5µM CHIR in TeSR-E6 for 1 hour. After 1 hour, cells were cultured with 1.1mL of TeSR-E6 with one of two growth factor combinations for 5 days, before being cultured in TeSR-E6 only to day 27. Media was changed every second day unless changing to a new media composition on a set protocol day. Samples were collected at the end of the four stages of differentiation as defined by changes in media composition, with the final organoids collected at day 27 of culture.

### Design and execution of kidney organoid single-cell transcriptional profiling

The kidney organoid protocol utilised was first presented in Takasato et al., 2015^15^ and then refined to use TESR-E6 minimal basal media^24,52^. The most commonly used kidney organoid protocol^7^, differentiation progresses from pluripotency through 4 stages; mesodermal induction (posterior primitive streak), IM commitment, nephron formation via a mesenchyme to epithelial transition and finally patterning and segmentation of nephrons within a renal stroma^7,15,23^. This study was designed to capture samples at all four stages of differentiation for the standard kidney organoid protocol while also examining the impact of variations in the duration of initial mesodermal patterning (3, 4 or 5 days of CHIR; 3c, 4c, 5c) and the impact of additional signaling pathway agonists (Wnt, GDF and FGF signaling) and antagonists (inhibition of RA and BMP) on final organoid cellular composition (Figure 1A). These growth factors were selected to interrogate the impact of rostro-caudal and mediolateral shifts in differentiation on both on-target and off-target cell type specification.

All experiments were performed simultaneously using the same iPSC line to minimise batch effects. Further, each unique condition was differentiated into three separate cultures, pooling the three replicates at the point of single-cell capture to include technical variability. Finally, each sample was labelled using hash-tag oligos^55^ (HTOs) for sample multiplexing (Figure 1B).

Following quality control, 150,957 single cells were processed for analysis. Integration, dimensional reduction, and diffusion mapping (Figure 1C) showed stages 1 and 2 to be nearly unique populations, while stages 3 and 4 had a clearer transition as organoids matured (Figure 1D).

### scRNA-seq capture including Hash-Tag Oligo sample barcoding

Kidney organoids were dissociated to single cell, incubated with Totalseq-A antibodies (Biolegend; Cat# 394601, 394603, 394605, 394607, 394609, 394611, 394613, 394615) for sample hashing each samples collection day, pooled and super-loaded onto the 10x Genomics Chromium Controller (10x Genomics) to capture single cells. Briefly, 3x technical replicates for each sample were dissociated using TrypLE for 3 minutes, pooled and diluted in TEsR-E6 and centrifuged at 300 × g for three minutes. The cell pellets were resuspended in 100 µL cold Fluorescence-Activated Cell Sorting (FACS) buffer (PBS with 2% fetal bovine serum). Then, 2 µL of Totalseq-A hashing antibody was added and the cells were gently pipetted to mix before incubating for 20 min on ice. Cells were then washed twice with FACS buffer by centrifuging at 300 × g for 5 min, discarding the supernatant and resuspending the cell pellet in 100 µL cold FACS buffer. Approximately 40,000 cells were loaded onto the Chromium Single Cell Chip B (10x Genomics; Catalog Number: PN-1000073) to capture 20,000 single cells with the Gel Bead Kit V3.0 (10x Genomics; Catalog Number: PN-1000075). GEM generation, barcoding, cDNA amplification, and library construction were performed according to the 10x Genomics Chromium User Guide (CG000183). Libraries were prepared with the Single Cell 3’ V3.0 Library and Gel Bead Kit (10x Genomics; Catalog Number: PN-1000075). The samples collected from stages 1, 2 and 3 occurred across six days, each capture multiplexed up to eight unique conditions and all were sequenced on the same run. The final collection of stage four cells occurred separately, multiplexing either 3 or 4 conditions to allow for greater cell numbers per condition for the more complex “mature” organoids. This design allowed us to maximise our capacity for cell capture per condition while minimising the potential batch effects that would impair downstream analysis.

13 unique scRNA captures were collected across the protocol. 7 captures were completed in the first 14 days of the protocol and were sequenced in the same batch, while the 6 captures at day 27 were sequenced in a second batch. HTO library sequencing was performed on the NextSeq 500/550 using a NextSeq 60 cycles kit (Illumina, 20416695) while the mRNA sequencing was performed on the NovaSeq 6000 using a NovaSeq S4 230 cycles kit (Illumina, 20363961).

### Data processing and quality control

Cell Ranger (v1.3.1) was used to process and aggregate raw data from each of the samples returning a count matrix. Quality control and analysis was performed in R using *Seurat*^56^ (v4.2.0) unless otherwise specified. Cite-seq-count (v1.4.3) was used to process HTO reads. *HTOdemux* was used to identify from the HTO profile cells as negative, singlet or doublets. For day 27 organoids, *HTOdemux* was run a second time on negatives. Cells with more than 2000 genes or identified as singlets using *HTOdemux* were kept while the remaining data was filtered out for downstream analysis. Data normalisation, scaling, variable gene identification and cell cycle phase identity regression were performed using the default implementation of *SCTransform*^57^ within *Seurat* using the parameters in Table 1. Clustering was performed using k-means clustering implemented within the *FindNeighbours* and *FindClusters* functions within *Seurat* using the parameters in Table 1. Integration of the 13 single cell captures was performed using *simspec*^36^ package (v0.0.0.9000) using the CSS algorithm executed using *cluster_sim_spectrum* utilizing default paramenters and sequencing batches as the grouping variable.

**Table 1.** Quantification of cell numbers for all classified cellular identities stratified by Stage of differentiation and duration of initial mesodermal patterning.

| Annotation | iPSC | Stage1 | Stage2 | Stage3 | Stage4 |
| --- | --- | --- | --- | --- | --- |
| iPSC | 2964 | 0 | 0 | 0 | 0 |
| iPSC-like | 0 | 22 | 0 | 0 | 0 |
| NMP | 0 | 81 | 0 | 0 | 0 |
| early_pPS | 0 | 3271 | 0 | 0 | 0 |
| late_pPS_1 | 0 | 2498 | 0 | 0 | 0 |
| late_pPS_2 | 0 | 3822 | 0 | 0 | 0 |
| pPSM | 0 | 0 | 2161 | 0 | 0 |
| IM-1 | 0 | 0 | 11037 | 0 | 0 |
| IM-2 | 0 | 0 | 6788 | 0 | 0 |
| Mesoderm-1 | 0 | 0 | 5909 | 0 | 0 |
| Mesoderm-2 | 0 | 0 | 6458 | 0 | 0 |
| Mesoderm-3 | 0 | 0 | 1530 | 0 | 0 |
| PM | 0 | 0 | 6308 | 0 | 0 |
| Neural-Prog | 0 | 0 | 4091 | 0 | 0 |
| IM-3 | 0 | 0 | 0 | 3405 | 0 |
| NPC-1 | 0 | 0 | 0 | 4045 | 0 |
| pNPC | 0 | 0 | 0 | 303 | 0 |
| Early_Nephron | 0 | 0 | 0 | 2053 | 0 |
| Early_Stroma | 0 | 0 | 0 | 753 | 0 |
| AdvMeso-1 | 0 | 0 | 0 | 5673 | 0 |
| AdvMeso-2 | 0 | 0 | 0 | 4445 | 0 |
| AdvMeso-3 | 0 | 0 | 0 | 2752 | 0 |
| Early_Neuron | 0 | 0 | 0 | 2053 | 0 |
| Neural_crest | 0 | 0 | 0 | 1786 | 0 |
| Myotome | 0 | 0 | 0 | 166 | 0 |
| UrEp | 0 | 0 | 0 | 2 | 0 |
| NPC-2 | 0 | 0 | 0 | 0 | 1444 |
| EN | 0 | 0 | 0 | 0 | 1659 |
| PEC | 0 | 0 | 0 | 0 | 2734 |
| Epod | 0 | 0 | 0 | 0 | 2412 |
| Pod | 0 | 0 | 0 | 0 | 1110 |
| EPT | 0 | 0 | 0 | 0 | 1720 |
| PT | 0 | 0 | 0 | 0 | 55 |
| EDT | 0 | 0 | 0 | 0 | 5902 |
| LOH | 0 | 0 | 0 | 0 | 211 |
| DT | 0 | 0 | 0 | 0 | 384 |
| UOS | 0 | 0 | 0 | 0 | 25 |
| Utip | 0 | 0 | 0 | 0 | 11 |
| UIS | 0 | 0 | 0 | 0 | 131 |
| SPC | 0 | 0 | 0 | 0 | 4 |
| MS | 0 | 0 | 0 | 0 | 721 |
| CS | 0 | 0 | 0 | 0 | 7025 |
| MesS | 0 | 0 | 0 | 0 | 6628 |
| Endo | 0 | 0 | 0 | 0 | 3 |
| Glia-Ependymal | 0 | 0 | 0 | 0 | 6834 |
| Glia-Schwann | 0 | 0 | 0 | 0 | 2374 |
| Glia-1 | 0 | 0 | 0 | 0 | 7929 |
| Glia-2 | 0 | 0 | 0 | 0 | 2011 |
| Glia-3 | 0 | 0 | 0 | 0 | 1157 |
| Myocyte | 0 | 0 | 0 | 0 | 834 |
| Satellite-cells | 0 | 0 | 0 | 0 | 3609 |
| Neuron | 0 | 0 | 0 | 0 | 2257 |
| NPC-like | 0 | 0 | 0 | 0 | 447 |

| Annotation | 3C | 4C | 5C |
| --- | --- | --- | --- |
| iPSC | 0 | 0 | 0 |
| iPSC-like | 12 | 4 | 6 |
| NMP | 22 | 28 | 31 |
| early_pPS | 2999 | 217 | 55 |
| late_pPS_1 | 9 | 1592 | 897 |
| late_pPS_2 | 27 | 1456 | 2339 |
| pPSM | 95 | 16 | 2050 |
| IM-1 | 2692 | 5144 | 3201 |
| IM-2 | 2295 | 2995 | 1498 |
| Mesoderm-1 | 1696 | 2029 | 2184 |
| Mesoderm-2 | 1920 | 1890 | 2648 |
| Mesoderm-3 | 514 | 524 | 492 |
| PM | 2178 | 2083 | 2047 |
| Neural-Prog | 1295 | 1823 | 973 |
| IM-3 | 1996 | 1123 | 286 |
| NPC-1 | 1429 | 1655 | 961 |
| pNPC | 144 | 55 | 104 |
| Early_Nephron | 2544 | 3218 | 2352 |
| Early_Stroma | 19 | 317 | 417 |
| AdvMeso-1 | 3393 | 1193 | 1087 |
| AdvMeso-2 | 416 | 1291 | 2738 |
| AdvMeso-3 | 1267 | 466 | 1019 |
| Early_Neuron | 921 | 3218 | 240 |
| Neural_crest | 721 | 765 | 300 |
| Myotome | 111 | 515 | 429 |
| UrEp | 0 | 1 | 1 |
| NPC-2 | 253 | 864 | 327 |
| EN | 311 | 864 | 484 |
| PEC | 576 | 960 | 1198 |
| Epod | 568 | 1030 | 814 |
| Pod | 108 | 258 | 744 |
| EPT | 227 | 412 | 1081 |
| PT | 3 | 15 | 37 |
| EDT | 1362 | 2235 | 2305 |
| LOH | 22 | 96 | 93 |
| DT | 109 | 167 | 108 |
| UOS | 11 | 9 | 5 |
| Utip | 3 | 8 | 0 |
| UIS | 83 | 35 | 13 |
| SPC | 0 | 0 | 4 |
| MS | 46 | 116 | 559 |
| CS | 835 | 2014 | 4176 |
| MesS | 803 | 2177 | 3684 |
| Endo | 2 | 1 | 0 |
| Glia-Ependymal | 5862 | 951 | 21 |
| Glia-Schwann | 812 | 919 | 643 |
| Glia-1 | 2492 | 2928 | 2509 |
| Glia-2 | 691 | 773 | 547 |
| Glia-3 | 296 | 61 | 800 |
| Myocyte | 259 | 234 | 341 |
| Satellite-cells | 1004 | 1680 | 925 |
| Neuron | 1151 | 395 | 711 |
| NPC-like | 48 | 114 | 255 |

### Use of R packages for reference-based cell classification

5 tools were used with data from each reference to further interrogate cellular identity within stages 1 and 2. *SCINA*^30^ uses the expression of gene sets within a data point to predict similarity to cell types, annotating cells above a particular level. *scAnnotatR*^33^ trains a classification model based on enriched gene sets to predict identity. *scClassify*^32^ generates cell-type hierarchies based on ensemble learning and sample size estimation. *SciBet*^31^ uses an entropy test to identify sets of genes enriched in a reference that provides significant association with cellular identity, using these to predict identity in a query. *scPred*^29^ uses machine learning to train classifiers for annotated references based on shared principal components and eigenvalues. *DevKidCC*^6^ utilises a set of pre-trained binary classifiers in a three-tier hierarchy of classification levels to robustly identify fetal kidney cell types in single cell data sets.

### Azimuth label transfer

A Seurat object was imported to the shiny application hosted at https://app.azimuth.hubmapconsortium.org/app/human-fetus after which the resulting files were downloaded for further analysis using R.

## Code and data availability

Code and data generated within this study will be available upon peer-review publication of this study.

